# Transgenic mice overexpressing Pitx2 in the atria develop tachycardia-bradycardia syndrome

**DOI:** 10.1101/2023.12.12.571383

**Authors:** Shunsuke Baba, Satoko Shinjo, Daiki Seya, Hiroki Bochimoto, Toru Akaike, Atsushi Nakano, Susumu Minamisawa

## Abstract

Sinoatrial node (SAN) dysfunction often accompanies supraventricular tachyarrhythmias such as atrial fibrillation (AF), which is referred to as tachycardia-bradycardia syndrome (TBS). Although there have been many studies on electrical remodeling in TBS, the regulatory mechanisms that cause electrical remodeling in the SAN and atrial muscles by chronic bradycardia or tachycardia have not yet been fully investigated. Here we hypothesized that Pitx2c, a transcription factor that plays a central role in the late aspects of left-right asymmetric morphogenesis, regulates an interrelationship between the SAN and the atrial muscles and is involved in TBS-like pathology. To test this hypothesis, we generated transgenic mice overexpressing *Pitx2c* specifically in the atria (OE mice). Although Pitx2c is normally expressed only in left atria, the expression levels of Pitx2c protein in the right atria were significantly increased to similar levels of those in the left atria of non-transgenic control mice (WT). We found that the heart rate of OE mice was significantly variable although the average heart rate was similar between WT and OE mice. Electrophysiological examination showed that OE mice exhibited prolonged SAN recovery time and higher AF inducibility. In addition, recording of the atrial monophasic action potential duration using a Langendorff perfusion system demonstrated shorter action potential duration in OE atria. Histological analysis revealed that SAN-specific ion channel HCN4-positive cells were hardly detected in the SAN of OE mice, along with ectopic expression in the right atria. Furthermore, transcription factors associated with sinus node formation were down-regulated in the right atria of OE mice. Therefore, SAN dysfunction by Pitx2 dysregulation predisposed OE mice to a TBS-like phenotype. We conclude that Pitx2c is a key regulator that defines SAN function in the atria.

## Introduction

The sinoatrial node (SAN) is essential for normal electrical function of the heart, allowing it to generate heartbeats and to adapt to changes in cardiac output and metabolic demand ^1^. The SAN, the primary pacemaker of the heart, is normally located at the junction of the right atrium (RA) and the superior vena cava (SVC), near the crista terminalis. Sick sinus syndrome (SSS) is caused by chronic SAN dysfunction and gives rise to profound sinus bradycardia and sinus pause (SP). Interestingly, approximately 50% of patients with SSS also have supraventricular tachyarrhythmias such as atrial fibrillation (AF) ^2^. This unusual combination of bradycardia and tachycardia has been known for more than 50 years and is referred to as tachycardia-bradycardia syndrome (TBS). In large population studies, the estimated hazard ratio for new onset AF was 4.2 in patients with SAN dysfunction^2^. One possible cause is that persistent bradycardia alters the electrical properties of the atrial muscle (electrical remodeling), making it more susceptible to tachycardia induction. On the other hand, it has been observed in clinical cases and animal studies that persistent atrial tachycardia, such as AF, can lead to SAN dysfunction. Moreover, clinical profiles of familial SSS probands with the mutation of HCN4 that is the SAN-specific ion channel ^3^ tended to exhibit AF with SSS ^4^. There have been many studies of electrical remodeling in TBS, including changes in ion channel expression and function.

However, the regulatory mechanisms that cause electrical remodeling in the SAN and atrial muscles by chronic bradycardia or tachycardia have not been fully investigated. We hypothesized, therefore, that there is an interrelationship between the SAN and the atrial muscles that might be strictly regulated at the transcriptional level. If this regulatory mechanism were to break down, a TBS-like phenotype could arise, in which disturbances in heart rhythm produce new disturbances in heart rhythm.

*Pitx2* is a paired-related homeobox gene that has been shown to play a central role in the late aspects of left-right asymmetric morphogenesis. *PITX2* was first identified as the gene responsible for Rieger’s syndrome and was recently reported to participate in the L-R asymmetrical pattern formation in body plan ^5^. The *Pitx2* locus produces five isoforms, designated *Pitx2a-e*, that arise through differential splicing and alternative promoter usage ^6^. In particular, *Pitx2c* is asymmetrically expressed in the left side of the developing heart ^7^. *Pitx2c* has been suggested to play a critical role in inhibiting the pacemaker gene program in the left atrium (LA), because *Pitx2c*-deficient mice exhibit ectopic automaticity in the LA ^8^. The lack of a *Pitx2c* gene showed a number of developmental defects, including the abnormal formation and function of SAN ^9^ ^10^, was susceptible to AF due to electrical remodeling^11^, and exhibited atrial tachycardia with irregular R–R intervals in adults ^12^. In line with these observations, genome-wide association studies (GWASs) showed that *PITX2* was a single variant near the 4q25 locus that conferred an over 60% increased risk of AF even in younger individuals ^13^. Furthermore, it has been reported that *PITX2c* mRNA expression was significantly higher in human atrial myocytes from chronic AF patients with an increase in the slow delayed rectifier K^+^ current density and a reduction in the voltage-gated Ca^2+^ current density ^14^. Putting these findings together, it is clear that proper expression of *Pitx2c* is important to maintain the electrical property of the atria. Therefore, we focused on Pitx2c to regulate the interrelationship between the SAN and the atrial muscles as a key molecule of TBS pathology. To test the hypothesis, we generated transgenic mice overexpressing *Pitx2c* specifically in the atria (OE mice) and found that these mice exhibited a TBS-like phenotype.

## Methods

### Genetically Modified Mice

We generated Pitx2-transgenic constructs using the pCAG-loxPSTOPloxP-ZsGreen vector (#51269, addgene, MA, USA). The transgenic construct is composed of a CAG promoter, two loxP sites flanking the STOP cassette, and mouse *Pitx2c* cDNA (NM_001042502.2) (Supplemental Figure 1A). The transgenic construct was injected into C57BL/6J mouse zygotes, and two mice positive for the transgene were obtained (founders), called B6-Tg (*CAG-LSL-Pitx2c*) transgenic mice. The founders were mated with wild-type (WT) C57BL/6J mice to obtain the F1 generation that was kept as a heterogeneous transgenic line. The presence of the transgene was confirmed by polymerase chain reaction (PCR).

The heterozygous *sarcolipin (SLN)* mutant mice (*SLN*^Cre/+^) that harbored the Cre recombinase cDNA and the defect of the *SLN* exon in one allele were described in detail elsewhere ^15^. In *SLN*^Cre/+^mice, Cre recombinase is exclusively expressed in the atria and the proximal regions of the junctional veins (SVC, intravena cava, and pulmonary veins) under the control of the endogenous *SLN* promoter. It should be noted that heterozygous SLN knockout mice have normal cardiac function without electrophysiological abnormality^16^. When a B6-Tg (*CAG-LSL-Pitx2c*) transgenic mouse was mated with a *SLN*^Cre/+^ mouse, we obtained four types of mice: wild type, B6-Tg (*CAG-LSL-Pitx2c*), *SLN*^Cre/+^, and B6-Tg (*CAG-LSL-Pitx2c*); *SLN*^Cre/+^ mice (Supplemental Figure 1B). We then examined the cardiac phenotypes of WT and B6-Tg (*CAG-LSL-Pitx2c*); *SLN*^Cre/+^(OE) mice in which the *Pitx2c* gene was considered to be selectively overexpressed in both right and left atria. All animal care and study protocols were approved by the Animal Ethics Committees of The Jikei University (2017-024C3), and the investigation conforms to the Guide for the Care and Use of Laboratory Animals published by the National Institutes of Health Guide for the Care and Use of Laboratory Animals (NIH Publication, revised 2011). Male and female mice of the same age were studied to ensure uniformity among each experimental subject. The mice were fed normal food and water that were freely accessed. Animals were housed in plexiglass cages and were kept in a room with controlled temperature (22 ± 3°C) and lighting (lights on from 7:00 AM–7:00 PM).

### Mouse genotyping

DNA for PCR screening was extracted from tail samples at 4 to 6 weeks after birth. Screening of Cre and *Pitx2* floxed alleles was routinely done as recommended by the manufacturer-specific primers (Supplemental Table 1). Cycling conditions for *Pitx2* and *SLN* were as follows: 1 min at 94°C, 35 cycles of 30 s at 94°C, 30 s at 58°C, and 30 s at 72°C, followed by a final extension step of 10 min at 72°C. PCR products were separated in standard agarose electrophoresis and classified according to the expected band size.

### Histology

For tissue preparation, whole heart samples were immobilized and fixed with 4% paraformaldehyde in 0.2 M phosphate buffer for one night, embedded in paraffin, sectioned longitudinally (4 µm), and stained with either hematoxylin-eosin (H&E) or Masson trichrome (MT) for interstitial fibrosis using standard protocols.

### Immunohistochemistry

For the detection of Pitx2c, the appendages of both atria were fixed in 10% formalin neutral buffer solution and then embedded in paraffin. Sections of 4 μm were mounted on slides. After deparaffination and rehydration, these sections were blocked in phosphate-buffered saline (PBS) containing 1.5% normal horse serum for 1 h at room temperature and incubated in primary antibody against Pitx2 (1:200, INNOVAGEN, Sweden) for 1 h at room temperature. Sections were washed and then incubated in secondary antibody (Alexa 488) for 1 h at room temperature and mounted using VECTASHIELD with DAPI (VECTOR Laboratories Ltd, UK) and coverslips.

For the detection of HCN4 and connexin43, whole heart samples were immobilized and fixed with 4% paraformaldehyde in 0.2 M phosphate buffer for 1 h, and soaked in 7.5% and 30% sucrose solution for 3 h. The whole heart with atria and supra vena cava was embedded in OCT compound (Sakura Finetek, Japan) and frozen by liquid nitrogen. Frozen serial sections (18 μm thickness) were cut along the frontal plane in a cryostat (Thermo Fisher Scientific, US) at -20℃. The cut sections were placed onto slides (MUTO PURE CHEMICALS CO Ltd, Japan) and stored at -80℃ until further use. Frozen serial sections were permeabilized 2 times with 0.2% Triton X-100 in high salt PBS for 5 min and then washed in high salt PBS. Next, sections were blocked in PBS containing 1.5% normal horse serum for 1 h at room temperature, washed in high salt PBS, and incubated in primary antibody of anti-HCN4 antibody (1:200, Alomone Labs, Israel) and anti connexin43 antibody (1:500, Sigma-Aldrich CO, US) for 3 nights at 4℃. Sections were then washed and incubated in secondary antibodies (Alexa Fluor 488, Alexa Fluor 594, Thermo Fisher Scientific, US) for 1 h at room temperature, and mounted using VECTASHIELD with DAPI and coverslips. Images were collected on an HS All-in-one Fluorescence Microscope (KEYENCE, Japan) using BZ-II software ^17^. Laser light wavelengths and filters appropriate for the secondary antibody fluorophores were used ^18^.

### Echocardiogram

Echocardiographic analysis of heart structure and function was performed at 8 weeks with surface electrocardiogram (ECG) using the Vevo 3100 device (FUJIFILM Visual Sonics Inc, Japan) with a 40 MHz high-frequency transducer (MX550D) as previously described ^19^. Mice were anesthetized with inhaled isoflurane (0.5–2%) delivered via nose cone while maintaining the heart rate over 400 bpm.

The left precordial hair was removed with hair removal cream. Body temperature was maintained with a heated imaging platform. In the parasternal short-axis projections, two-dimensional images were recorded at the mid-papillary muscle level and pulse wave doppler images were recorded at the right ventricle outflow tract level with guided M-mode recordings. Heart rate and fractional shortening were measured and averaged from each image for at least 3 beats. Additionally, in order to assess the pulmonary hypertension resulting from morphological abnormality of the pulmonary vein, the pulmonary acceleration time (PAT) and pulmonary ejection time (PET) were measured and averaged from each image for at least 3 beats and the PAT to PET ratio was calculated. No arrhythmia was observed in any mice during the recording.

### Telemetry Electrocardiography

To assess heart rate in conscious and unrestrained animals, a radio transmitter (Data Sciences International, US) was implanted in each mouse at 10 to 12 weeks. Mice were anesthetized with 1 to 2% inhaled isoflurane and the abdominal and chest hair was removed with hair removal cream. Next, the pedal reflex in the mice was checked to confirm successful anesthesia prior to operation. An abdominal midline incision was made on the ventral surface. Two other smaller incisions were made in the pectoral region for suturing of the leads. The positive lead of the transmitter was tunneled subcutaneously to the left anterior chest wall above the apex of the heart, and the negative lead was placed to the right shoulder. This configuration approximates body surface ECG of leadⅡ (lead II).

Subcutaneously, these leads were directed cranially from the abdominal incision and sutured into the pectoral muscles ^20^. The 3.5 g wireless radiofrequency transmitter was then inserted into the subcutaneous or abdominal cavity. On the day after the operation, to allow for adequate recovery from surgery, each unrestrained mouse, housed in a separate and isolated cage far away from the stimulation of other animals, was placed on top of a radio receiver (Data Sciences International, US) in a room with regulated temperature conditions (22 ± 1℃) and synchronized to a light–dark schedule of 12 h to 12 h (lights on from 7:00 AM–7:00 PM) to conserve the circadian rhythm of each animal.

The ECG signals were continuously monitored and recorded for 48 h. Mice were free to eat and drink during the recording ^20^. ECG signal data were processed using a computer-assisted data acquisition system (Data Sciences International, US) that yielded a time series of the ECG waveform, and values for electrography analysis for every epoch as prescribed ^21^ ^22^.

### ECG waveform analysis

The RR interval, PR interval, QRS duration, and QT interval were measured in lead II. Since the QT interval covaries with the RR interval, we calculated a rate-corrected QT interval (QTc) using the formula modified for mice as prescribed ^23^. All waves in 24 h were manually checked by the same researcher. To assess the circadian variation in heart rate and the variability of the heart rate, each mouse was measured for the standard deviation of heart rate every hour and the standard deviation was averaged over 24 h; RR durations per every epoch were analyzed every hour. To calculate the averaged PR interval, QRS interval, and QTc interval, 10 epochs were sampled every hour from a 24 h monitoring session ^24^.

In addition, arrhythmia events were counted in each mouse. Premature atrial contraction (PAC) was defined as premature P waves with premature QRS complexes that made the premature beat less than two normal cycles. SP was defined as an R-R interval 50% greater than the average R-R interval from the telemetric data in each group ^17^.

### Electrophysiological examination

Animals underwent catheter-based intracardiac stimulation and recordings with the body surface ECG. Mice were anesthetized using 1 to 2% inhaled isoflurane to maintain a heart rate between 300 to 400 bpm during the experiment and the pedal reflex was checked to confirm successful anesthesia prior to examination. Next, atropine sulfate (4 μg/g) (NiproES pharma, Japan) and propranolol hydrochloride (1 μg/g) (AstraZeneca, UK) were sequentially injected into the abdominal cavity to measure the SAN activity without the effect of the autonomic regulation of sinus function ^25, 26^ . Then, subcutaneous needle electrodes placed immediately under the skin were used to obtain leadⅡ using a polygraph system (NIHONKODEN, Japan) and the Ponemah Physiology Platform software (Data Sciences International, US). For the intravenous stimulation and electrograms, we inserted a 1.6 F octapolar catheter (Millar, US) connected to ADI BioAmp and PowerLab apparatus with an isolator (NIHONKODEN, Japan) to stimulate or with Lab Chart7 Software (AD Instruments, New Zealand) to record the intravenous electrograms from a right internal jugular vein. A vertical skin cut-down over the right jugular vein was performed, and the vein was isolated from connective tissue. A small transverse nick was made in the vein, and the catheter was advanced through the nick and jugular vein to the RA using intracardiac electrograms as guides. Correct catheter placement was confirmed by having a maximal atrial signal in the distal lead and being able to stimulate the atrium adequately following the stimulation of the atrium from the intracardiac catheter in the distal lead ^22^. After the trial stimulation to make sure the catheter was at the correct position and to check the capture threshold amplitudes, the atrium was paced at 1.5 times the capture threshold amplitudes. Sinus node recovery time (SNRT) was measured three times after delivering an atrial pacing train for 30 s with a cycle length of 130 to 180 ms depending on the baseline heart rate. SNRT was defined as the duration between the last pacing stimuli and the onset of the first spontaneous P wave of the body surface leadⅡ. Rate-corrected SNRT (CSNRT) was calculated by subtracting the baseline heart rate from maximum SNRT among three trials ^27^.

Next, the inducibility of AF was analyzed by applying a 15 s burst using an automated stimulator as described previously ^28, 29^. A series of burst pacing was repeated three times after stabilization. AF was defined as a period of rapid irregular atrial rhythm lasting at least 2 s. The duration of AF was defined as the interval between the beginning of irregular atrial rhythm following the burst pacing and the onset of the first normal sinus beat no longer than 20 min ^30^. These experiments were performed 30 to 40 min after the injection of the drugs to ensure that the drugs were effective, as shown in the experiment of the pharmacological autonomic nervous blockade (not shown), and to avoid the effect of anesthesia.

### Quantitative real-time polymerase chain reaction

We performed quantitative real-time polymerase chain reaction (q-RTPCR) as previously described ^16^. Briefly, total RNA was extracted from atrium tissues using Direct-zol RNA MicroPrep kit (Zymo Research, US) and reverse-transcribed into cDNA using Superscript RT Master Mix (Takara, Japan) as recommended by the manufacturer. We performed q-RTPCR using TB Green Premix Ex Taq Ⅱ (Takara, Japan) with specific primers (Supplemental Table 1), and TaKaRa Thermal Cycler Dice (TaKaRa, Japan) as recommended by the manufacturer. Target mRNA expression was normalized to *Gapdh* mRNA expression as an internal reference. The relative expression quantity 2^(−△Ct)^ was used to calculate the differences among groups.

### Western blot analysis

Three or four snap-frozen atrial tissues were prepared and homogenized in cold RIPA lysis buffer containing PMSF, protease inhibitors, and sodium orthovanadate using a RIPA lysis buffer system Santa Cruz Biotechnology, US). The preparation was centrifuged at 15000 rpm at 4℃ for 5 min. The protein concentration was quantified using the TaKaRa BCA Protein Assay Kit (TaKaRa, Japan), and 2-mercaptoethanol (Nacalai Tesque, Japan) was added to the protein lysate to the appropriate concentration. Proteins (30 μg/lane) were separated by electrophoresis on SDS-PAGE gels (BioRad, US) and blotted onto PVDF membranes (MERCK MILLIPORE, Germany). The membrane was blocked in 5% milk in TBST (0.05% Tween 20 in TBS) and incubated with appropriate primary antibodies overnight at 4℃. The anti-HCN4 antibody (Sigma-Aldrich, US) was used at 1:200 dilution in TBST, the anti-PITX2 ABC antibody (INNOVAGEN, Sweden) at 1:500 dilution, and the anti-GAPDH antibody (Cell Signaling Technology, US) at 1:1000 dilution. Membranes were then washed three times in TBST and incubated with appropriate horseradish peroxidase–conjugated secondary antibodies at 1:5000 dilution in TBST (Cell Signaling Technology, US) for 1 h at room temperature. We quantified the expression of each of these proteins at the predicted molecular weights according to the information from the supplier. Signal acquisition was performed using EZ-Capture MG (ATTO, Japan). The intensities of the proteins were measured by ImageJ (National Institutes of Health, US) software and were normalized to the intensity of endogenous GAPDH. The bands density was calculated for each sample and used for the comparison between groups. For another antibody, the membrane was stripped using stripping buffer (Thermo Fisher Scientific, US) and washed with TBST after detection.

### Preparation of Hearts for Langendorff Perfusion

We administered intraperitoneal heparin (0.1 ml by intraperitoneal administration) to mice 10 min before heart dissection. After mice were anesthetized and had no response to pain stimuli, their hearts were isolated quickly and connected to a Langendorff perfusion setup as described previously ^16, 31^.

After an initial period of 10 min to allow the preparation to stabilize, the heart was perfused at a constant perfusion pressure of 80 mmHg with a 37°C warm oxygenated HEPES-Tyrode solution containing 137 mM NaCl, 5.4 mM KCl, 0.5 mM MgCl_2_ 0.33mM NaH_2_PO_4_, 5 mM HEPES, and 5 mM glucose; pH adjusted to 7.4.

### Recording of Atrial Monophasic Action Potential During Langendorff Perfusion

Atrial monophasic action potentials (MAPs) were recorded from the atrium of isolated hearts using a microprobe. This miniaturized contact MAP probe could be held at an atrial epicardial site as required with a heart holder including a stimulating electrode, and enabled stable recordings from the atrial epicardial heart surface.

After confirming that the probe was in the proper position and recording the characteristic MAP waveform of the atrium as prescribed ^32^, atrial MAP was recorded in 10 s without pacing and with pacing by 8 Hz, 10 Hz, and 12 Hz. The action potential duration (APD) was determined at different repolarization levels (APD20, APD60, and APD90) with full repolarization defined as 100% and the tip of the MAP upstroke as 0%. Time of maximum upstroke velocity (dV/dtmax) was defined as time Each APD was measured and averaged from each heart at 3 beats.

### Statistics

All data are expressed as means ± standard error of the mean (SEM). Statistical analyses were performed after assessing for normal distribution using unpaired *t-*tests with Welch’s correction for comparison of two groups. Detection of the genotype effect, atrial laterality effects, and interaction between genotype and atrial laterality were tested with a repeated measures two-way analysis of variance (ANOVA). Values of *p* < 0.05 were considered statistically significant.

## Results

### Mice with the atrial cardiomyocyte–specific overexpression of the Pitx2c gene normally survived into adulthood without any apparent cardiac morphology

We generated transgenic mice which overexpress *Pitx2c* specifically in the atria by crossing SLN^cre/+^ mice and TG (LSL-Pitx2c) mice. OE mice were born with more than the expected Mendelian ratio (Supplemental Figure 1B), indicating that they do not have embryonic lethality. OE mice did not exhibit an abnormal appearance and could survive into adulthood with normal behavior, physical activity, and fertility. The ratios of atrial and ventricular weights to body weight were not different between WT and OE mice (Table 1).

**Figure 1.**
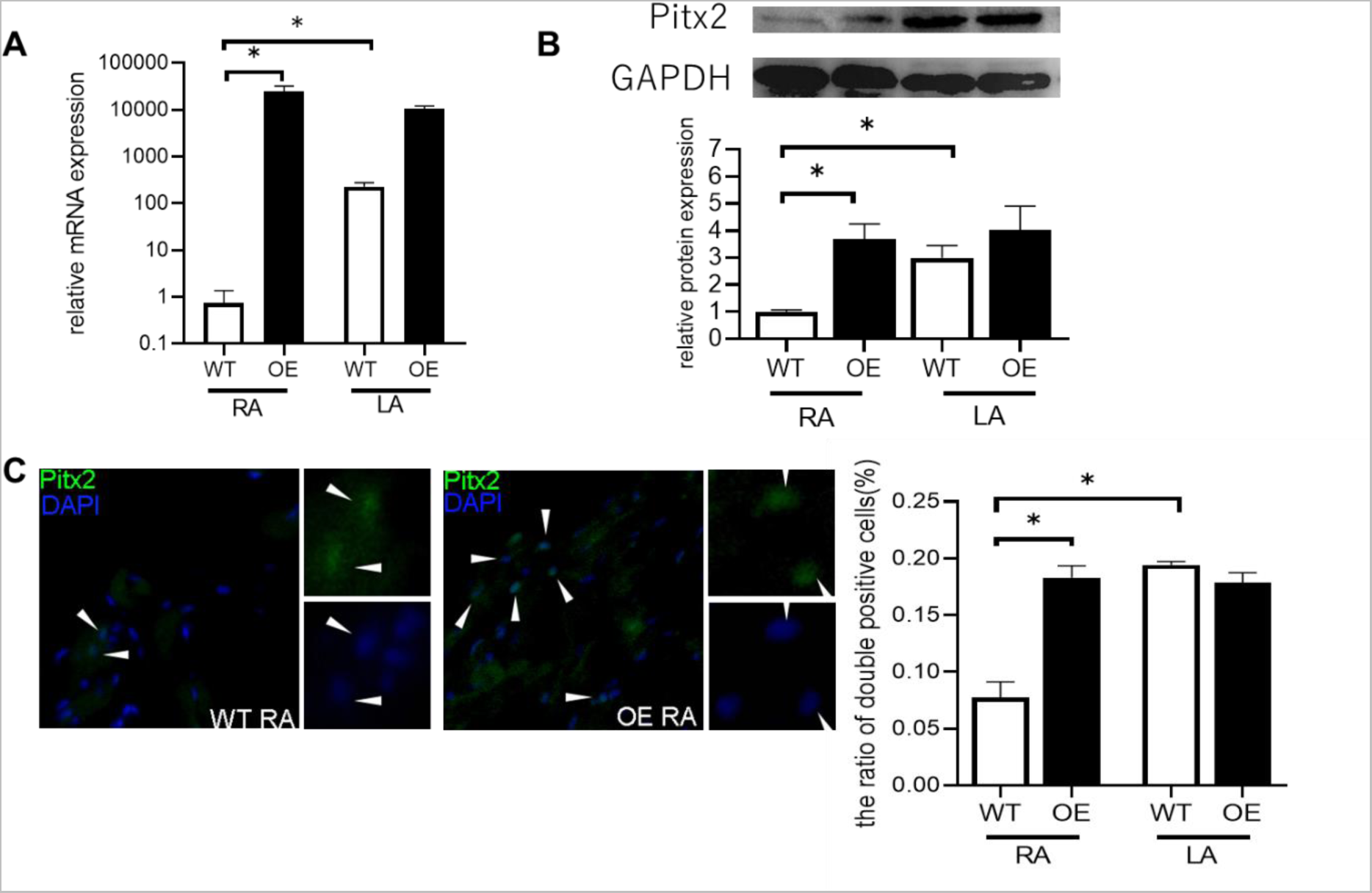
The expression of Pitx2c in the atria of WT and OE mice. A, RTPCR showing that the Pitx2c isoform in the right atrium of OE mice increased significantly. The data are the mean ± standard error of the mean (SEM) (n = 4 per group, \**p* < 0.05). B, Western blot showing that Pitx2 in the right atrium of OE mice increased significantly. The data are the mean ± SEM (n = 8 per group, \**p* < 0.05) from four independent experiments. C, The figure with the arrow indicates double-positive cells (Pitx2: green and DAPI: blue) and the graph with the averaged ratio of double-positive cell number in three fields of view under the microscope in each mouse. The data are mean ± SEM (n = 5 per group, \**p* < 0.05).

**Table 1.** Parameters of each group tc. The table shows the parameters for the ratio of atrial and ventricular weights to body weight in each group at 10 to 13 weeks, echocardiographic and electrocardiac evaluation at 8 weeks. The data are the mean ± standard error of the mean (n = 8 per group).

|  | **WT** | **OE** |
| --- | --- | --- |
| Age, days | 102.5 ± 5.3 | 105.0 ± 5.2 |
| BW, mg | 26.4 ± 1.8 | 23.9 ± 1.8 |
| LA/BW, × 10<sup>-2</sup> | 1.53 ± 0.10 | 1.25 ± 0.04 |
| RA/BW, × 10<sup>-2</sup> | 1.78 ± 0.13 | 1.56 ± 0.12 |
| LV/BW, × 10<sup>-2</sup> | 34.2 ± 1.19 | 41.0 ± 0.04 |
| RV/BW, × 10<sup>-2</sup> | 10.7 ± 0.66 | 11.5 ± 1.15 |
| <u>echocardiography</u> |  |  |
| HR, bpm | 433 ± 7.18 | 407 ± 23.9 |
| LVPWs, mm | 1.02 ± 0.06 | 0.84 ± 0.05 |
| LVPWd, mm | 0.76 ± 0.05 | 0.64 ± 0.08 |
| LVIDs, mm | 2.73 ± 0.17 | 2.91 ± 0.12 |
| LVIDd, mm | 3.81 ± 0.16 | 3.95 ± 0.11 |
| IVSs, mm | 1.05 ± 0.08 | 0.94 ± 0.04 |
| IVSd, mm | 0.65 ± 0.09 | 0.68 ± 0.05 |
| FS, % | 28.4 ± 2.46 | 26.6 ± 1.39 |
| P valve, mm | 1.41 ± 0.07 | 1.38 ± 0.06 |
| PA VTI, mm | 21.0 ± 2.31 | 18.7 ± 1.30 |
| CO, ml/min | 13.9 ± 1.65 | 11.3 ± 1.22 |
| PAT, ms | 17.6 ± 1.08 | 18.7 ± 1.28 |
| PET, ms | 61.6 ± 2.36 | 67.1 ± 2.85 |
| PAT/PET | 0.29 ± 0.02 | 0.28 ± 0.02 |
| <u>electrocardiography</u> |  |  |
| PR, ms | 34.8 ± 0.91 | 34.0 ± 1.41 |
| QRS, ms | 13.1 ± 0.20 | 13.2 ± 0.53 |
| QTc, ms | 40.2 ± 0.67 | 43.3 ± 1.45 |

As previously reported, we also found that the expression levels of *Pitx2c* mRNA were significantly higher in the LA than in the RA in WT mice and that *Pitx2c* was the predominant isoform when compared to *Pitx2a* and *Pitx2b* isoforms in the mouse atria, especially in the RA (Figure 1A, Supplemental Figure 2) ^11^. RTPCR analyses showed that the expression levels of *Pitx2c* mRNA were significantly increased in the RA of OE mice compared to those of WT mice and there was no statistical difference in the expression levels of *Pitx2c* mRNA between the RA and LA of OE mice. At the protein level, western blot analysis for total Pitx2 showed that the expression levels of Pitx2 protein in the RA of OE mice were significantly increased to similar levels of the LA of WT and OE mice (Figure 1B). Furthermore, the number of Pitx2 and DAPI double-positive cells was significantly increased in the RA of OE mice (Figure 1C). These data indicated that we successfully obtained mice with atrial cardiomyocyte–specific overexpression of the *Pitx2c* gene.

**Figure 2.**
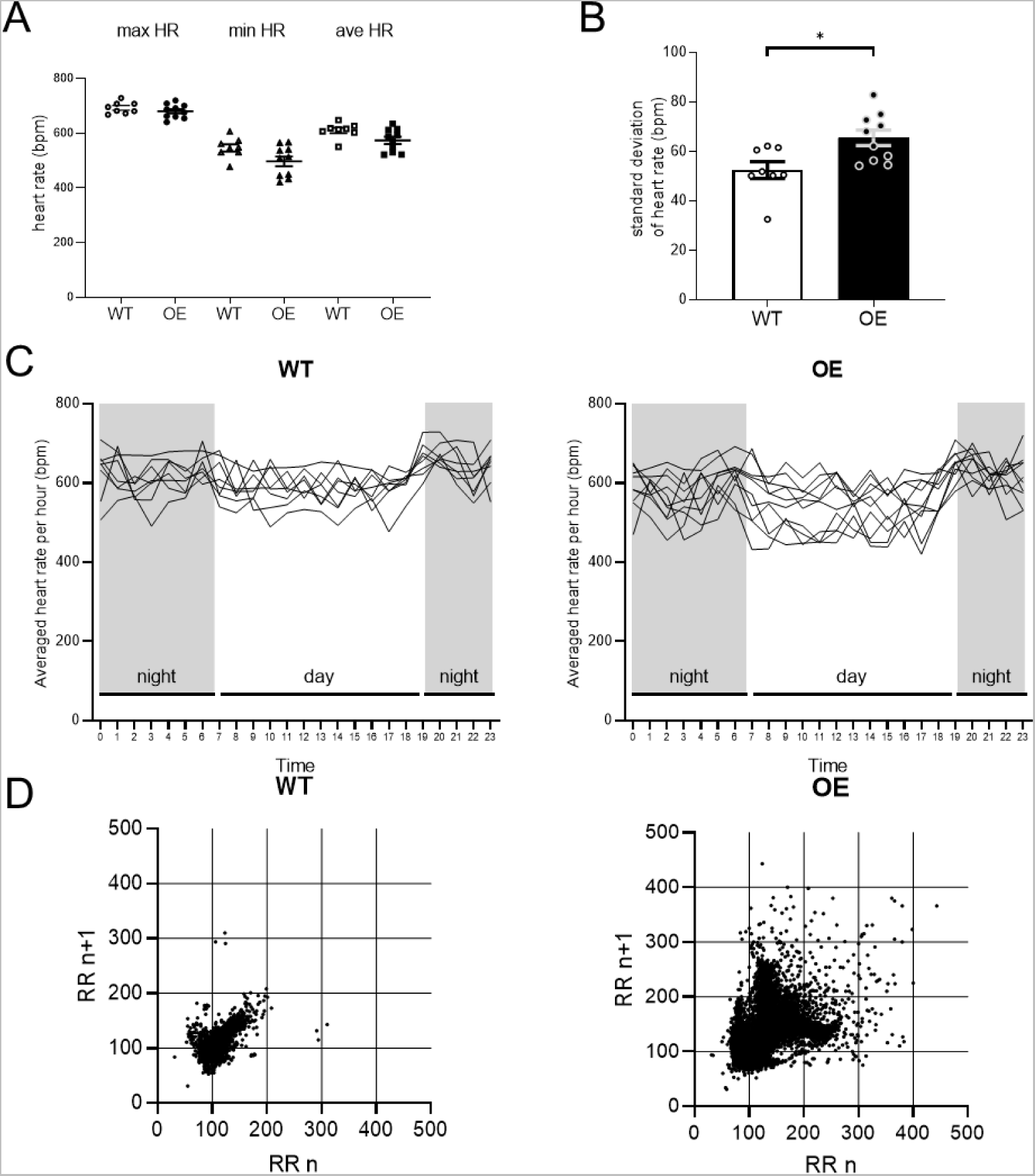
Variability of heart rate in WT and OE mice. A, B, and C, The telemetric data show that the variability of the heart rate increased significantly in OE mice. There was no significant difference in the max/min/ave heart rate of each mouse for 24 h. The data are the mean ± standard error of the mean (WT n = 8, OE n = 10, \**p* < 0.05) D, Lorenz plots show the consecutive RR intervals in each group

The global morphology of the atria showed the usual appearance and arrangement of the RA and LA in OE mice (Supplemental Figure 3), suggesting that the atrial morphology was similar between the WT and OE mice. In addition, the ratio of collagen fibrosis to myocardium was similar between WT and OE mice using MT staining.

Echocardiographic evaluation also indicated that the parameters of cardiac function were similar in both WT and OE mice at 8 weeks old (Table 1). In addition, we did not find any structural abnormality in OE hearts using echocardiography.

### Heart rate variability significantly increased and atrial arrhythmias were frequently observed in OE mice

A spot ECG recording at echocardiogram examination showed no difference in PR interval, QRS interval, and QT interval between WT and OE mice (Table 1) and no abnormal waves such as SP and ectopic beats. We then continuously measured ECG using telemetry monitoring. We found no significant difference in the maximum HR, minimum HR, and averaged HR between OE and WT mice (Figure 2A). However, the standard deviation of the HR significantly increased in OE mice (Figure 2B). In line with this, the 24 h ECG recording showed that the oscillation range of the average heart rate per hour was wider in OE mice than in WT mice (Figure 2C). Lorenz plots analysis of the RR intervals distinguished by telemetric data also indicated that the heart rate variability significantly increased in OE mice (Figure 2D).

In the telemetric data, we often found a number of atrial arrhythmias, including PAC and SP in OE mice (Figure 3A, 3B). Moreover, OE mice frequently showed waves of AF and tachycardia-bradycardia (Figure 3C, 3D). These data indicated that the variability of the heart rate significantly increased in OE mice and that atrial arrhythmias could contribute to the variability.

**Figure 3.**
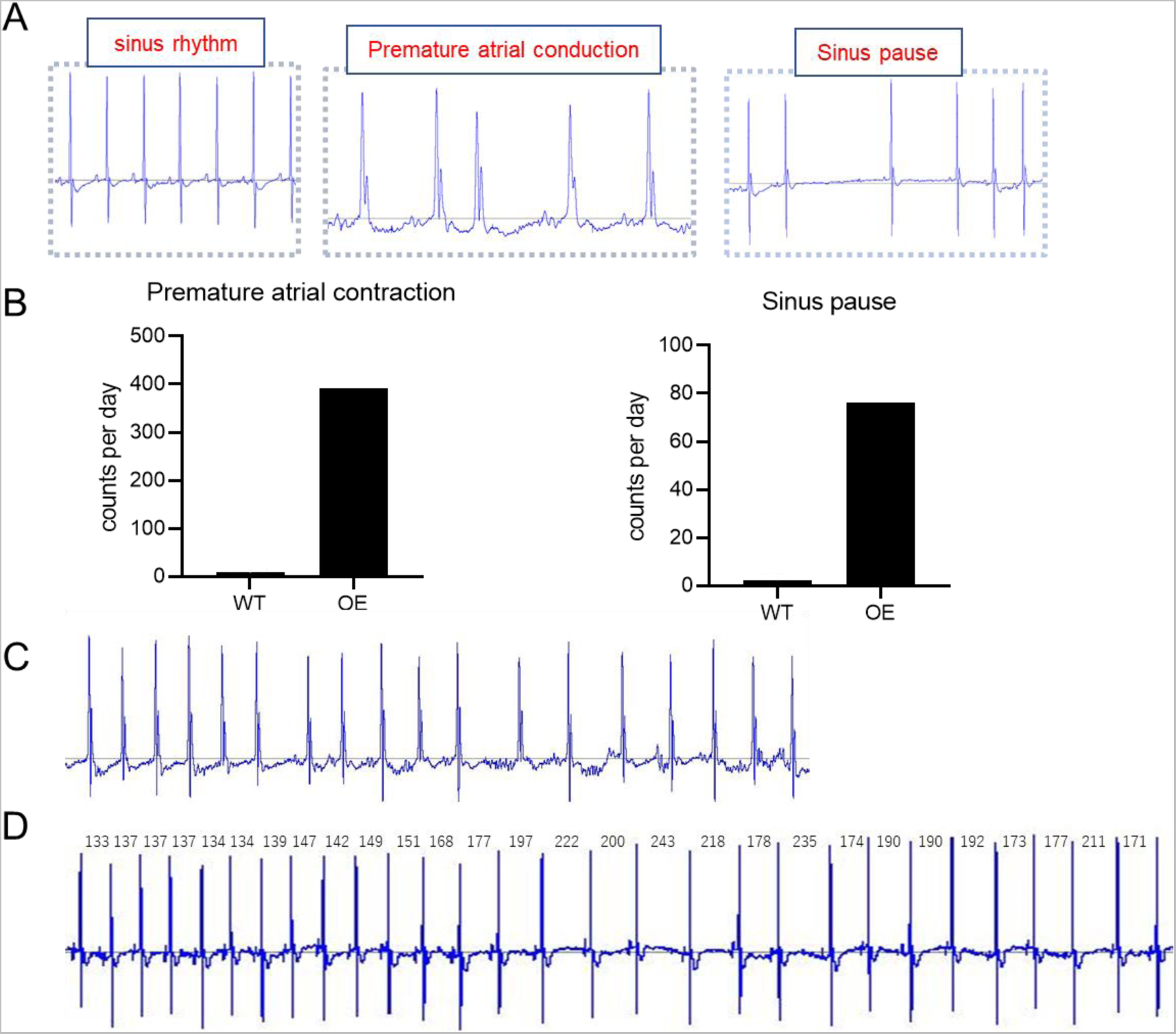
Atrial arrhythmias were frequently observed in OE mice. A, The telemetric data show representative examples of ECG waves for sinus rhythm and arrhythmias B, The number of arrhythmia events (PAC and SP) increased in OE mice. C and D, The telemetric data recorded the waves of AF and tachycardia-bradycardia. The number above the wave shows each RR interval length.

### OE mice exhibited sinus node dysfunction and higher AF inducibility

We evaluated the SAN function of *in vivo* mice using the overdrive suppression test. The maximal SNRT of OE mice was longer than that of WT mice. Furthermore, the CSNRT of OE mice was significantly longer than that of WT mice (Figure 4, Supplemental Table 2), indicating that the SAN function was impaired in OE mice. Intracardiac electrical burst stimulation induced reproducible AF more frequently in OE mice than in WT mice (OE: 51.9%, WT: 5.6%), suggesting that the OE mice were more susceptible to AF than WT mice.

**Figure 4.**
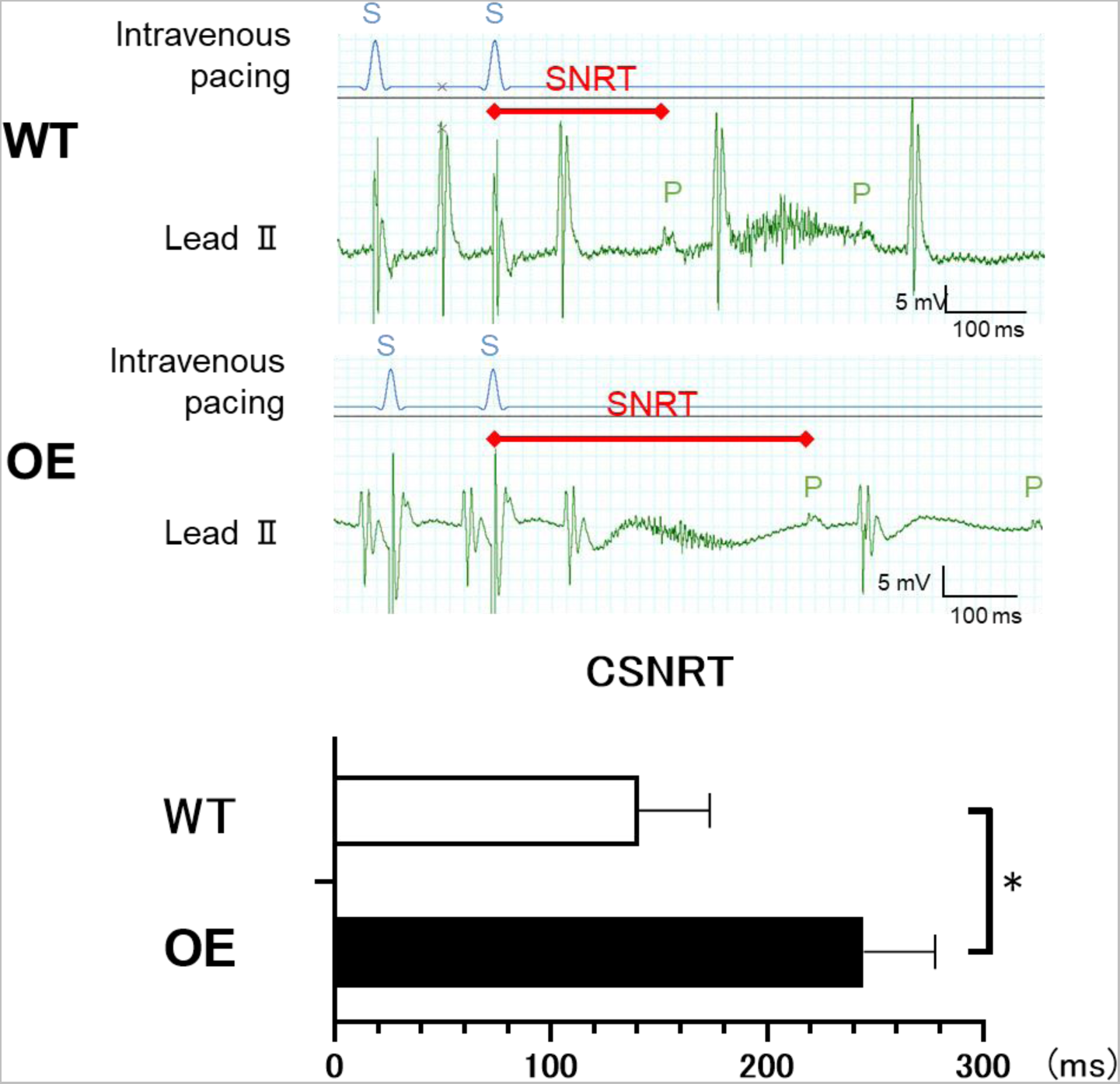
Overdrive suppression test demonstrated SAN dysfunction in OE mice. Representative records of the overdrive suppression test. SNRT of OE mice was longer than that of WT mice. (WT n = 6, OE n = 9, \**p* < 0.05) S: stimulus wave of the intravenous pacing; P: P wave of lead Ⅱ on the surface; SNRT was defined as the interval between the last stimulus in the pacing train and the onset of the first spontaneous atrial excitation. The rate-corrected sinus node recovery time (CSNRT) was defined as SNRT minus basic sinus cycle length.

### HCN4-positive cells were decreased in the SAN and HCN4 was ectopically expressed in the RA of OE mice

To identify whether the SAN is formed normally in OE mice, we examined the SAN by immunohistochemical HCN4 staining ^17, 18^. The area of the SAN which was identified as the HCN4-positive and Cx43-negative population was suppressed in OE mice compared with WT mice at the junction of the SVC and the RA. In contrast, HCN4-positive and Cx43-negative cells were ectopically expressed in the RA of OE mice (Figure 6).

**Figure 5.**
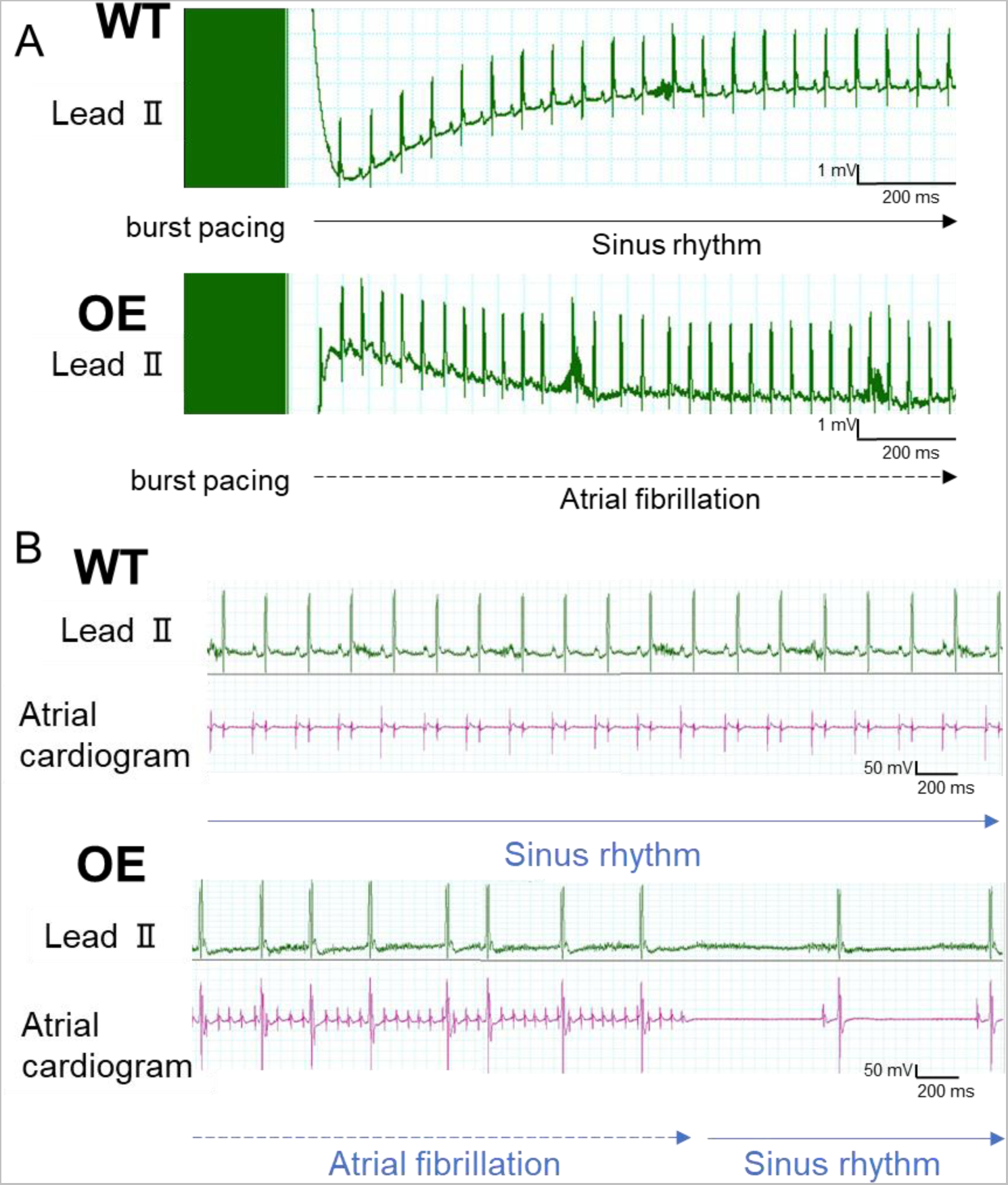
Intracardiac electrical stimulation studies and incidence of reproducible pacing-induced AF. A and B, A burst pacing protocol was used to induce AF. Simultaneous surface ECG (A) and intracardiac electrograms (B) revealed irregular R wave to R-wave intervals in OE mice after burst pacing, which are suggestive of AF with an irregularly irregular ventricular response.

**Figure 6.**
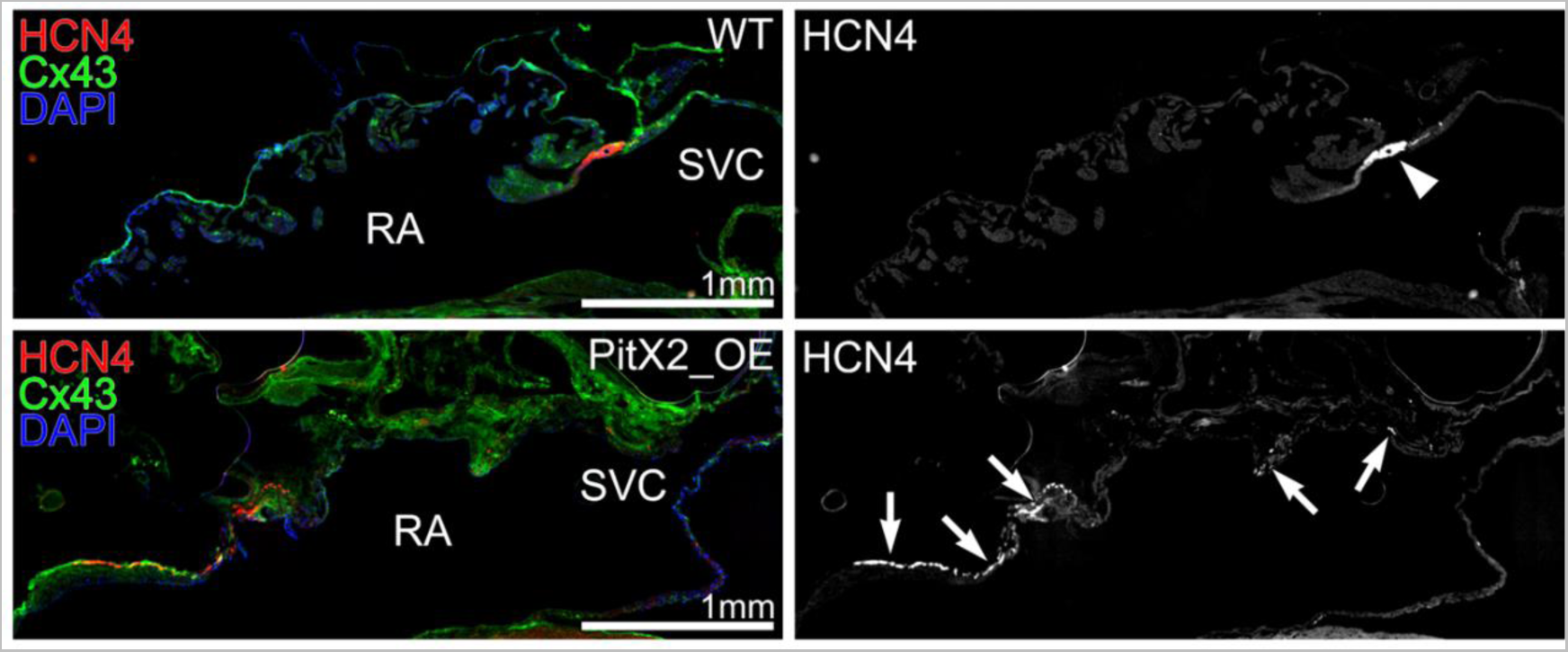
Immunohistochemical staining of RA and SVC for HCN4 and Cx43. Upper figure of WT mice showed the area of HCN4-positive and Cx43-negative population expressed at the junction of the SVC and the RA. The lower figure of OE mice showed that this area was ectopically expressed in the RA.

### Transcription factors related to SAN formation were down-regulated in the RA of OE mice

Since *Pitx2c* overexpression in the RA impaired the SAN formation and function as described above, we examined the transcription factors related to the left-right asymmetric morphogenesis and electrical remodeling of the SAN. *Tbx3*, *Tbx5*, and *Shox2* mRNAs, which were relevant to the specification of the SAN, were significantly down-regulated in OE mice (Supplemental Figure 4A). In contrast, the expression levels of *NKX2.5* mRNA were comparable between WT and OE atria.

### Action potential duration was shorter in OE atria with reduced expression of Ca^2+^ regulator genes

The monophasic action potential duration (APD) was shorter in the RA of OE mice (Table 2, Supplemental Figure 5). The expression levels of *HCN4*, *Cx40, Cx43,* S*cn5a, Cacna1c, Serca2, Ryr2, Pln*, and *Csq2* mRNAs were down-regulated in OE mice, and those of *Kcne1* mRNA were up-regulated in OE mice. In particular, *Serca2* and *Cacna1c*, which are known as intracellular calcium regulators, were significantly reduced in the RA of OE mice (Supplemental Figure 4B).

**Table 2.**
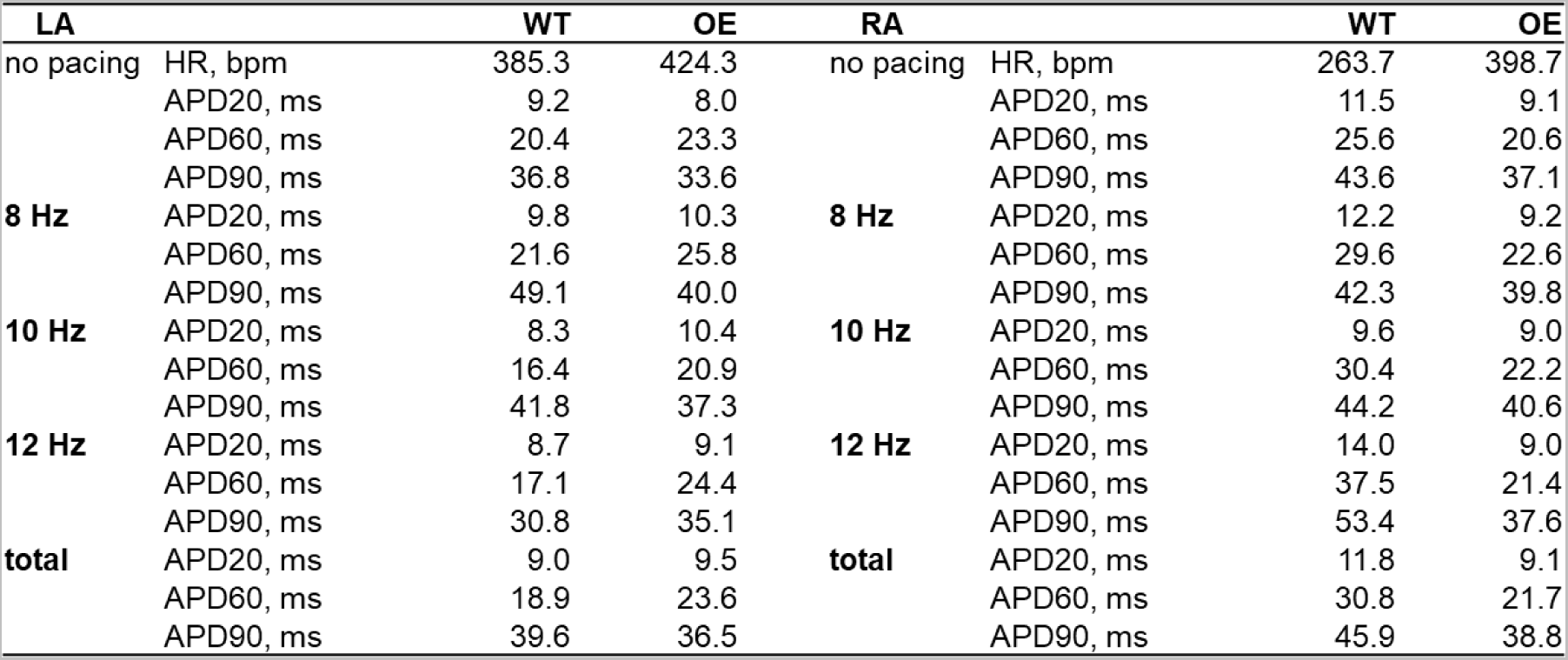
Comparison of atrial MAP of RA in OE and WT. APD at different repolarization levels (APD20, APD60, and APD90) without pacing and with pacing by 8 Hz, 10 Hz, and 12 Hz. The data are the mean ± standard error of the mean (n = 3 per group).

## Discussion

In the present study, we demonstrated that transgenic mice atrium-specific overexpressing *Pitx2c* exhibited SAN dysfunction and are prone to AF, which resembles a TBS-like phenotype. Previous studies using genetic screening and genetically modified animal models have elucidated the electrophysiological mechanisms of TBS ^33^. Namely, mutations of the ion channels underlying the SAN function are known to cause TBS-like phenotypes. For example, Duhme et al. demonstrated that mutation in the HCN4 gene (HCN4-K530N) is associated with familial TBS and persistent AF ^34^ and Ziyadeh-Isleem et al. showed that a truncating SCN5A mutation causes SSS and early onset of AF ^35^. However, little was known about the regulatory mechanisms by which electrical remodeling, including these ion channel changes, occurs in the SAN and atrial muscles as a result of chronic bradycardia or persistent tachycardia. Here, our results suggest that *Pitx2c* plays a key role in regulating the interrelationship between the SAN and the atrial muscles and that up-regulation of *Pitx2c* in the RA may be the first step in causing electrical remodeling via rhythm disturbances in the SAN and atrial muscles. Atrium-specific *Pitx2c* overexpressing mice would be an excellent animal model of TBS to investigate how rhythm disturbances promote electrical remodeling and generate other types of rhythm disturbances.

From an embryological perspective, the transcription factor cascades and signaling pathways that determine the formation and characterization of SAN are being studied to provide a complete picture. Pitx2c is thought to be a negative regulator of SAN development, because Pitx2c controls the asymmetric left-sided morphogenesis of the venous poles and Pitx2c-deficient fetuses in mice form the SAN not only in the RA, but also in the LA ^36^. However, whether forced expression of Pitx2c at the right-sided venous pole and the RA inhibits the SAN-specific gene program and directly suppresses the SAN formation has not yet been investigated *in vivo*. Therefore, to the best of our knowledge, this is the first *in vivo* evidence demonstrating that the presence of Pitx2c in the right-sided venous pole suppressed SAN formation, indicating that the absence of Pitx2 expression in the RA is critical for SAN formation during development. It is known that *Pitx2c* is an upstream transcription factor that directly or indirectly inhibits the expression of the genes involved in the SAN-specific gene program such as *Shox2*, *Tbx3*, and *Tbx5* ^37^. In our study, q-RTPCR showed that *Shox2*, *Tbx3*, and *Tbx5* were significantly down-regulated in the RA of OE mice. Even though the data were not obtained from SAN tissues, our data also supported that Pitx2c works as an inhibitory transcription factor of the SAN-specific gene program.

Many studies have demonstrated the relation between Pitx2 and AF. Previous experiments with genetically engineered mice have shown that a decrease in Pitx2c promotes electrical remodeling of the atria and predisposes mice to AF ^11, 12^. Interestingly, Pérez-Hernández et al. reported that the expression of *Pitx2c* mRNA was significantly higher in human atrial myocytes from chronic AF patients with an increase in the slow delayed rectifier K^+^ current density and a reduction in the voltage-gated Ca^2+^ current density ^14^. This data was recently confirmed by a simulation model study^38^. In the present study, we showed for the first time that overexpression of Pitx2c in the RA also induced susceptibility to AF *in vivo*. Taken together, it is suggested that the proper expression level of *Pitx2c* is important to maintain the stable electrical property of the atria. Although the exact molecular mechanism of how overexpression of Pitx2c in the RA causes arrhythmogenesis was not revealed in the present study, the ectopic expression of HCN4 and the shortened APD of the RA are very likely responsible for Pitx2c-mediated arrhythmogenesis. *Hcn4* is the isoform of hyperpolarization-activated cyclic nucleotide–gated channels and is mainly responsible for the slow diastolic depolarization of pacemaker cells. HCN4 is expressed exclusively in the cardiac conduction system, particularly in the SAN. When HCN4 is ectopically expressed in the working myocardium like OE mice, the areas expressing HCN4 would exhibit ectopic automaticity that is likely to induce AF. In the present study, considering the shape of the P waves in the telemetry electrocardiogram and electrophysiological study in OE mice, the locus of the ectopic beats was considered to be in the RA. Therefore, the spots where HCN4 was ectopically expressed in the RA of OE mice is likely to be the origin of arrhythmogenesis. Similarly, ectopic automaticity that induced AF with electrical remodeling was found in the LA of Pitx2c heterozygous-deficient mice in which HCN4 was ectopically expressed in the LA ^8^. For persistence of AF, the presence of ectopic automaticity is not sufficient; electrical remodeling in the atria is required for reentry to be established. In this process, shortening of the APD at the atria is a common phenomenon observed in AF, as observed in OE mice. A previous report showed that Ptx2c up-regulated the slow delayed rectifier K^+^ current and down-regulated the Na^+^ current, inward rectifier K^+^ current, and L-type Ca2^+^ current from AF promoting gene-regulatory networks ^38–41^. In our study, q-RTPCR showed that S*cn5a, Cx40, Cx43,* and *Cacna1c* were down-regulated, and *Kcne1* was up-regulated at the transcriptional levels in OE mice. *Kcne1* generates the slowly activating K^+^ current and *Cacna1c* encodes the L-type high voltage-gated calcium channel mediating the influx of calcium ions into the cell. Up-regulation of *Kcne1* and down-regulation of *Cacna1c* would contribute to shortening of the APD. *Scn5a* encodes a cardiac voltage–dependent sodium channel and *Cx40/43* are gap junction proteins that allow cardiac tissue to function as an electrically continuous syncytium. Down-regulation of *Scn5a* and *Cx40/43* might contribute to slowing atrial conduction in OE mice. Moreover, we also found down-regulation of *Serca2, Ryr2, Pln,* and *Csq2*, which are intracellular calcium regulator genes that would alter calcium cycling in OE mice. Future studies are needed to clarify whether there are changes in membrane currents or changes in intracellular calcium dynamics.

It should be noted that the OE mice used in this study overexpress pitx2c in an atrium-specific manner, but there were no obvious morphological abnormalities in the atria, including left-right symmetry. Since the expression of sarcolipin starts from embryonic day 10.5, the transgene for Pitx2c is also supposed to be expressed from embryonic day 10.5. In this time window, the asymmetric morphology of the atria is nearly complete, but the formation of the SAN is completed around embryonic day 13.5 ^42^. The atrial morphology is normal in OE mice, because the cre recombinase is expressed and functional after the completion of atrial morphogenesis but before the SAN is fully formed ^43^.

Finally, it is of note that the present results are attributed to the effects of Pitx2 overexpression in the atria from early embryonic life ^37, 38^. This experimental design does not allow us to determine whether, for some reason, Pitx2c expression increases in the RA in adulthood, after the SAN is already properly formed, and whether the increase in Pitx2c affects the function of the SAN. Therefore, it is necessary to create inducible Pitx2c overexpression mice or other means to investigate this important issue in future studies ^8, 9^.

## Acknowledgments

We thank Drs. Motoji Sawabe and Yurie Soejima at the Graduate School of Medical and Dental Sciences Biomedical Sciences and Engineering Division of Biomedical Laboratory Sciences Department of Molecular Pathology, Tokyo Medical and Dental University for critical discussion and comments about immunostaining.

## Non-standard Abbreviations and Acronyms

LA: left atrium;
RA: right atrium;
LV: left ventricular;
RV: right ventricular;
BW: body weight
HR: heart rate;
FS: fraction shortening;
PAT-PET: ratio of pulmonary acceleration time to pulmonary ejection time as an estimate of pulmonary hypertension
maxHR/minHR/aveHR: maximum/minimum/average heart rate in 24 h.
VHR: variability of heart rate
PAC: premature atrial conduction,
SP: sinus pause
SNRT: sinus node recovery;
CSNRT: rate-corrected sinus node recovery time
BHR, bpm: basic heart rate in beats per minute

---

**Supplemental Figure 1.**
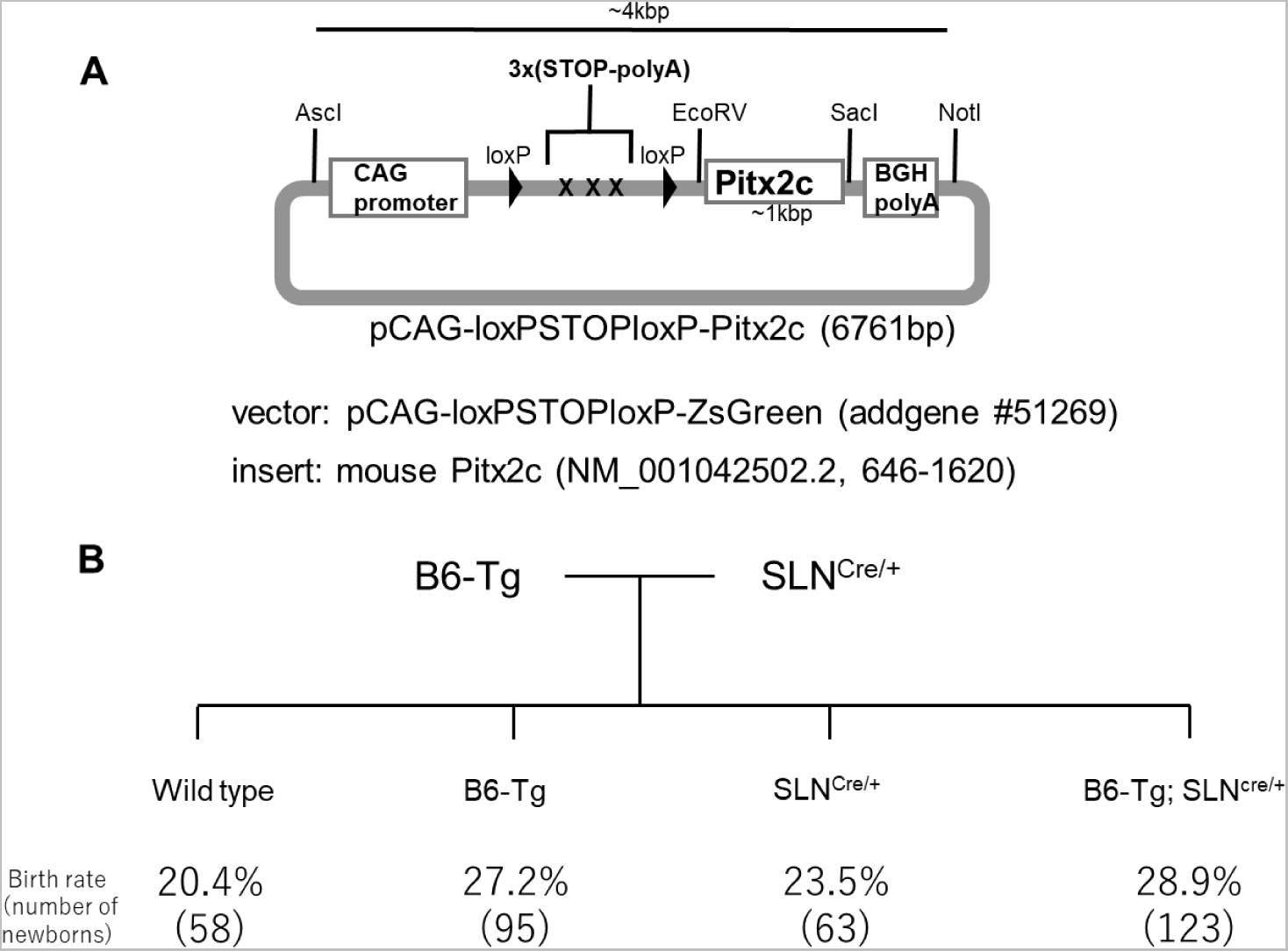
Transgenic construct and genotypes. A, We generated the genetic construct for the conditional overexpression of Pitx2c mice. B, We obtained four types of mice, which were born at more than the expected Mendelian ratio.

**Supplemental Figure 2.**
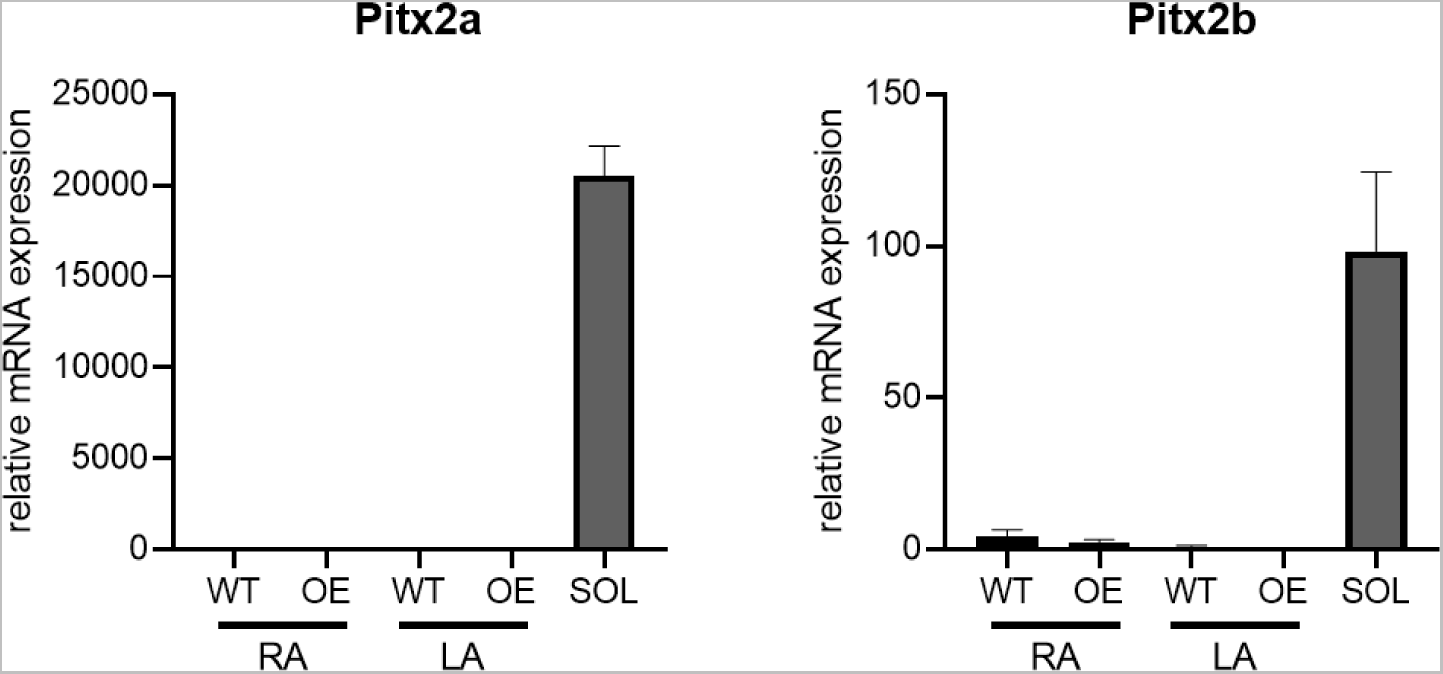
mRNA expression of Pitx2a and Pitx2b. RTPCR shows that the Pitx2a and Pitx2b isoform in the mouse heart is almost less expressed than the soleus as the positive control. The data are the mean ± standard error of the mean (n = 4 per group, #*p* < 0.05).

**Supplemental Figure 3.**
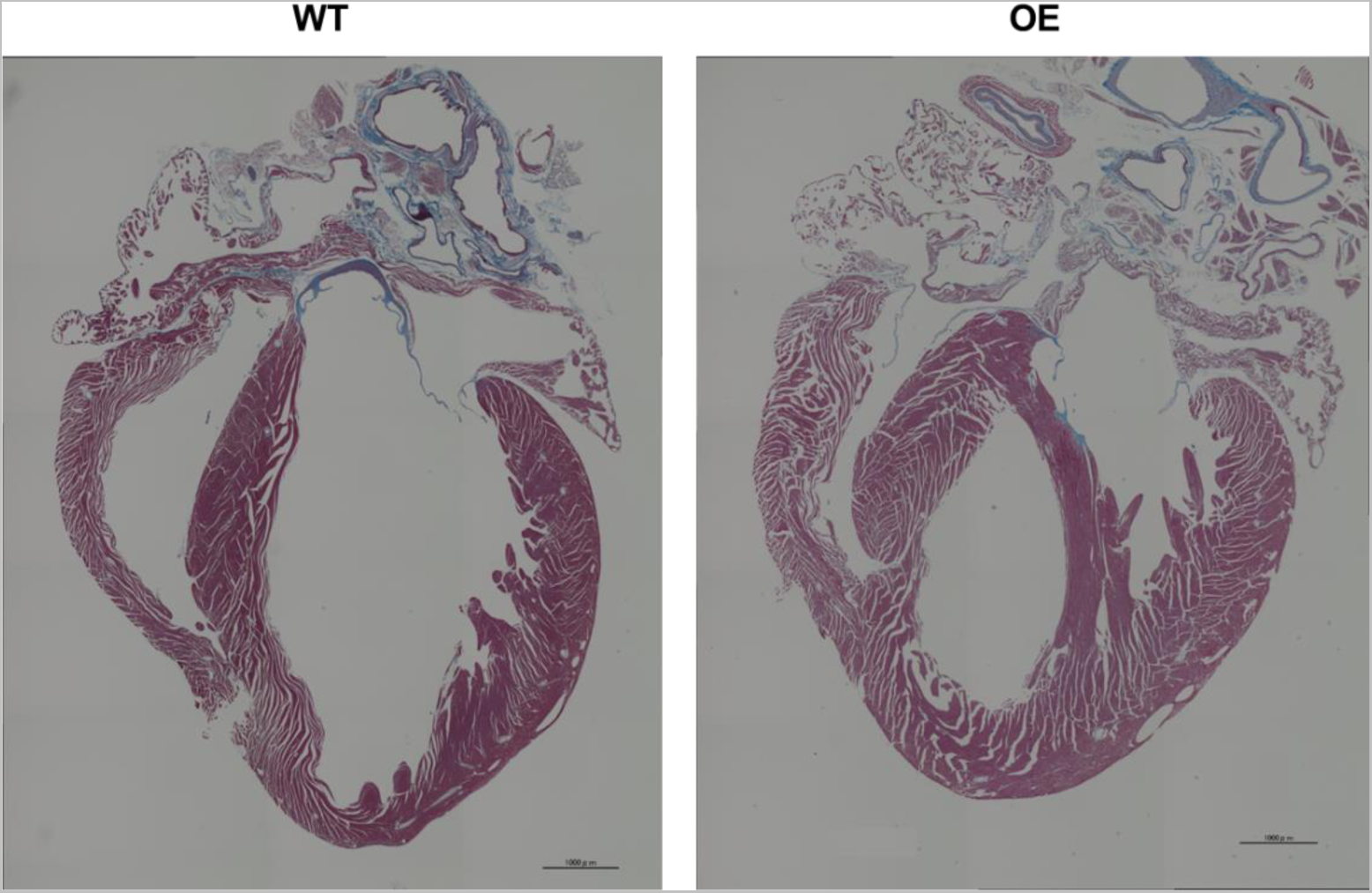
Histological analysis of the heart in WT and OE mice. These figures of MT staining of the whole heart show no difference in morphology and fibrosis between WT and OE mice.

**Supplemental Figure 4.**
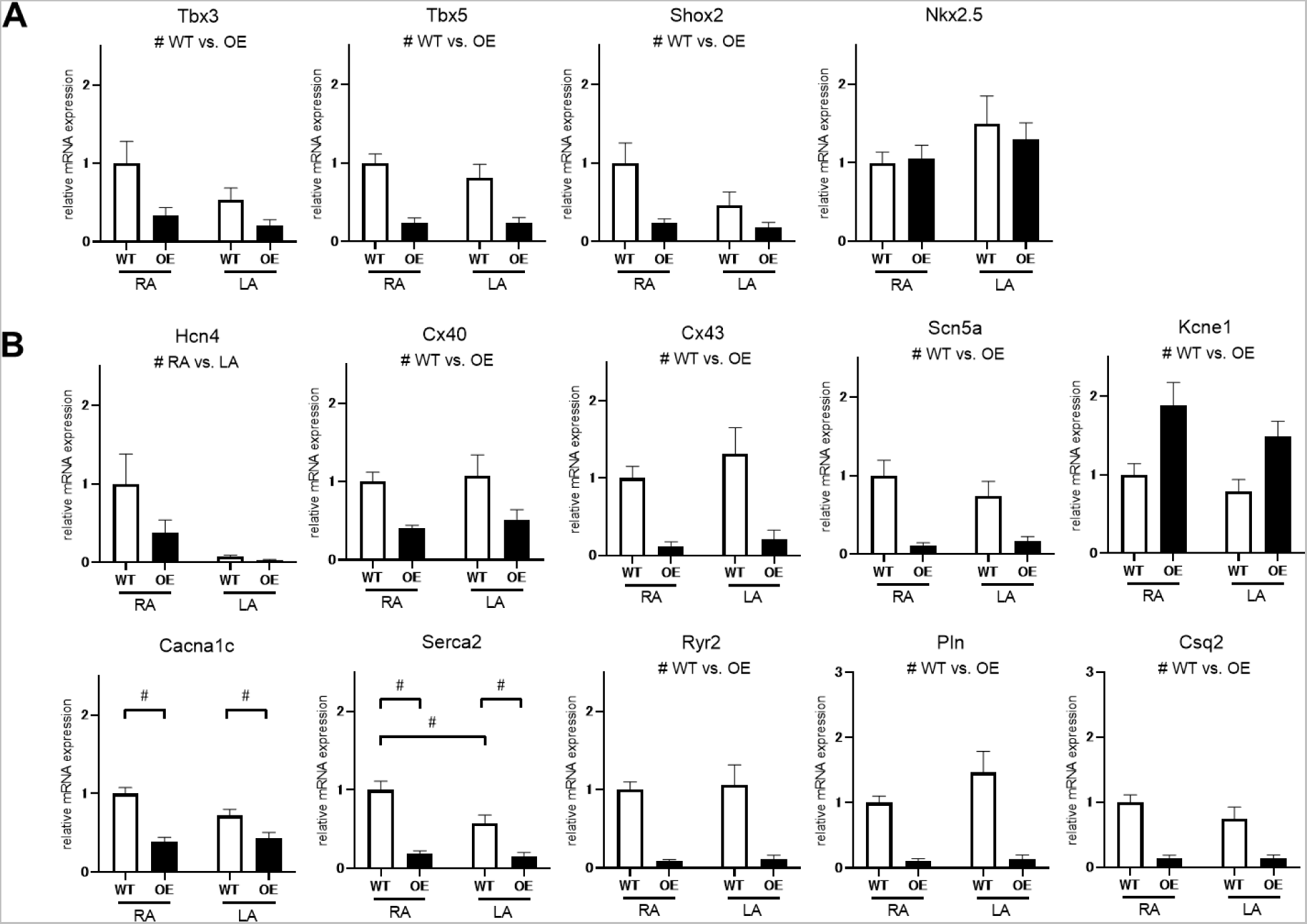
mRNA expression (A. transcription factor, B. not transcription factor) related to left-right asymmetric morphogenesis and the inducibility of AF. In OE mice, mRNAs were significantly changed. In particular, mRNAs (*Cacna1c*, *Serca2*) related to Ca handling which promoted AF were reduced in the RA of OE mice. The data are the mean ± standard error of the mean (n = 8 per group, #*p* < 0.05)

**Supplemental Figure 5.**
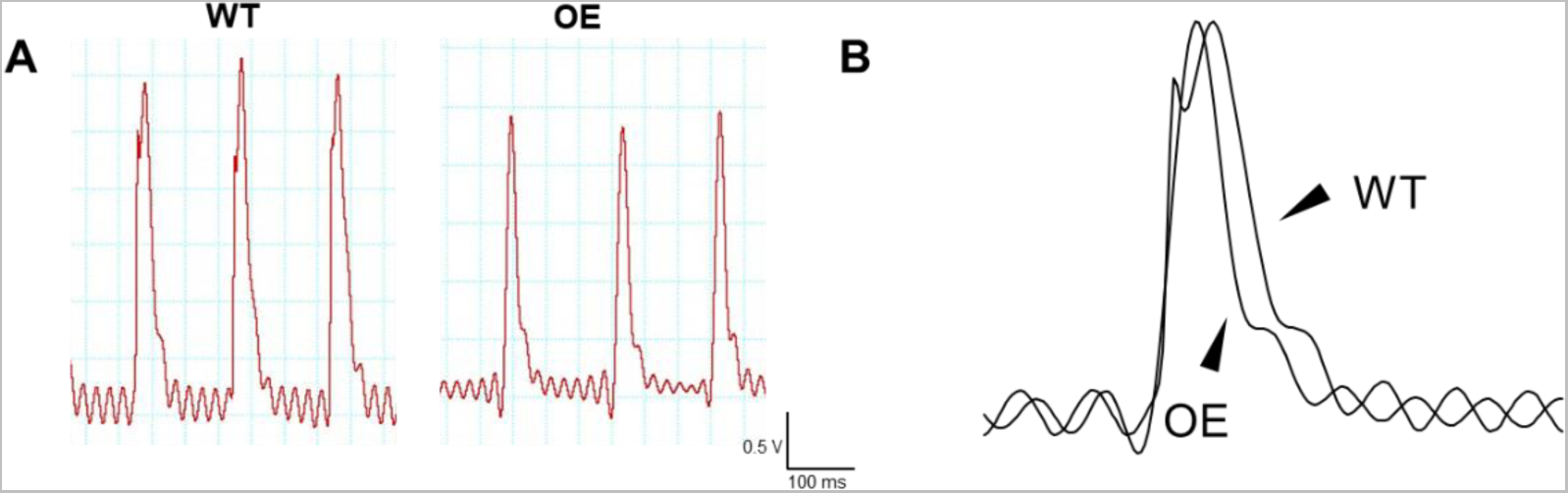
Comparison of atrial MAP of RA in OE and WT mice. A, Atrial MAP was recorded without pacing and with pacing by 8 Hz, 10 Hz, and 12 Hz. B, Overlay of recordings in the MAP about RA of OE mice and RA of WT mice.

**Supplemental Table 1.**
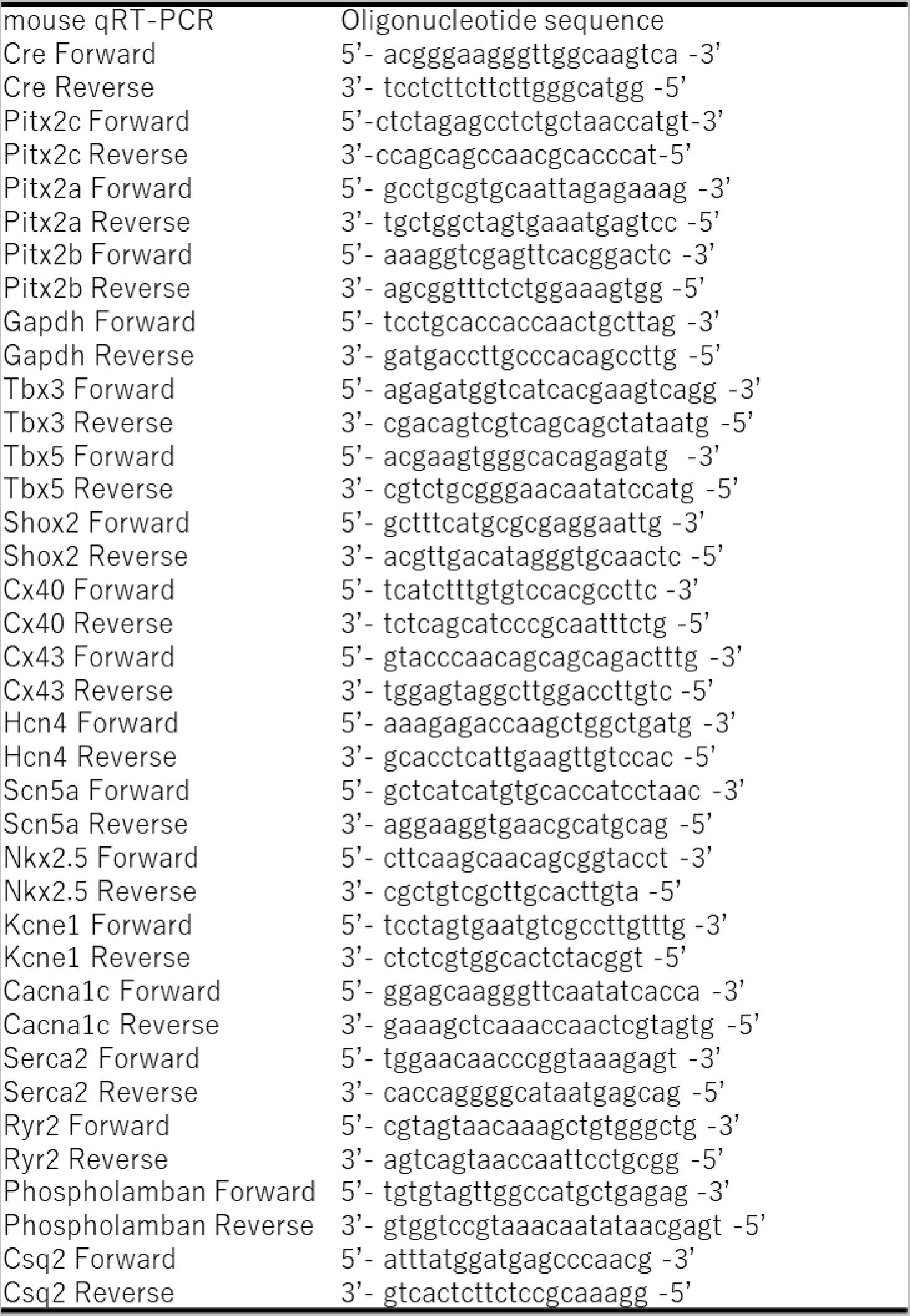
Primers which were used for polymerase chain reaction screening of the DNA.

**Supplemental Table 2.** Parameters for the overdrive suppression test. The data are the mean ± standard error of the mean (n = 8 per group). max SNRT: maximum sinus node recovery time of three trials; duration and amplitude: the duration and amplitude of the stimulation for 30 s. Total experimental duration: the experimental time from the anesthesia to the end.

|  | WT | OE |
| --- | --- | --- |
| max SNRT, ms | 358 ± 36.9 | 437 ± 32.4 |
| BHR, bpm | 280 ± 14.6 | 313 ± 8.10 |
| Cycle length, ms | 162 ± 4.00 | 156 ± 2.77 |
| amplitude, mV | 0.108 ± 0.0117 | 0.11 ± 0.0152 |
| total experimental duration, min | 31 ± 3.1 | 32 ± 1.8 |

